# Evaluating Reference-Independent Pipelines for the Detection of Spreading Organisms in Metagenomic Datasets

**DOI:** 10.64898/2026.05.03.722517

**Authors:** N.S. Popov, V.V. Panova, M. Molchanova, S.A. Gurov, A.N. Lukashev, E.N. Ilina, A.I. Manolov

## Abstract

The emergence of unidentified pathogens, or “Disease X,” poses a significant threat to global health, necessitating the development of proactive surveillance strategies for the wildlife and human virosphere. Since novel viruses often lack universal genetic markers or known homologs, this study evaluates four reference-independent computational pipelines: coverage-based, k-mer-based, nucleotide clustering, and Large Language Model (LLM)-based designed to detect spreading organisms by comparing distinct metagenomic datasets. Using a real-world pandemic dataset of human nasopharyngeal RNA-seq runs and a semi-synthetic dataset enriched with divergent Egovirales sequences, we measured the sensitivity, selectivity, and computational efficiency of each approach. The coverage-based method proved most robust, consistently achieving 100% genome coverage of SARS-CoV-2 and maintaining high selectivity even at low viral concentrations, though it required extensive computational resources (20 days of CPU time for 2B reads). In contrast, the k-mer-based approach offered a tenfold reduction in execution time and high selectivity but was sensitive to data depletion, failing to detect targets at very low abundances. The clustering-based pipeline performed effectively at moderate concentrations but suffered from sequence fragmentation in sparse data, while the LLM-based method (using ViraLM), despite its efficiency, exhibited critically low selectivity due to current latent space partitioning limitations. These results demonstrate that while k-mer and LLM-based tools provide rapid screening capabilities, the coverage-based approach remains the most reliable for sensitive pathogen discovery. Ultimately, these reference-independent workflows are essential for illuminating metagenomic “dark matter” and establishing early warning systems for emerging infectious diseases

## Introduction

The emergence of “Disease X”—a hypothetical, unidentified pathogen—poses a severe threat to global pandemic preparedness [1] [2]. The source of disease X may be wildlife, where it may spread cryptically [3]. Even more, it might spread in the human population without symptoms similar to HIV/AIDS. Because these pathogens often circulate cryptically in wildlife, proactive characterization of the wildlife virosphere is essential [4]. Since viruses lack a universal genetic marker, high-resolution metagenomic next-generation sequencing (mNGS) is the necessary tool for mapping viral diversity and assessing cross-species transmission risks.

The most obvious way to identify a new circulating agent is to compare two sets of samples obtained from similar sources using similar methods. Comparison by detecting the organism in initially negative samples or detection mutations in known organisms.

Sequence-dependent methods enable the identification of novel viral lineages through phylogenetic analysis of conserved regions, as shown by the discovery of a potential new HIV-1 subtype [5] and the SIVbkm strain of Simian Immunodeficiency Virus in Lophocebus aterrimus [6], both identified via genomic clustering with known primate lentiviruses.

In contrast, sequence-independent methods require no prior knowledge about the organism of interest. Non-computational approaches include suppression subtractive hybridization (SSH) and representational difference analysis (RDA) [7]. For example, SSH-based cDNA subtraction of infected/uninfected mouse liver RNA led to the identification and sequencing of a novel murine coronavirus (MHV-MI) [8]. As well, RDA has been successfully applied for the primary identification of a variety of organisms, including such viruses as TTV [9] and HHV-8 [10].

The shift from specialized laboratory protocols to high-throughput metagenomic workflows scales the discovery process by standardizing the experimental phase and transferring the identification of novel organisms into the computational domain through the analysis of raw sequence data.

New organisms are discovered via co-abundance profiling, which groups genetic sequences that appear at correlated concentrations across different samples. By cross-assembling viral metagenomes from 12 individuals, researchers identified contigs with highly correlated coverage depths, allowing them to reconstruct the 97 kbp circular genome of crAssphage. This widespread bacteriophage had remained hidden among unclassified sequences because its proteins lacked matches in existing sequence databases [11].

Ponsero et al. (2023) demonstrated that k-mer-based de novo methods reliably replicate taxonomic structures without reference databases, provided researchers use quantitative metrics, k-mer lengths ≥ 20 bp, and sufficient sequencing depth [12]. Similarly, using short k-mer spectra (5≤k≤11) allows for rapid beta-diversity analysis of shotgun metagenomic data on human gut metagenomes, enabling the detection of uncharacterized agents like crAssphage that alignment-based methods often miss [13]. Mathematical models further prove that k-mer differences directly reflect nucleotide diversity, allowing population parameters to be estimated directly from raw data [14]. Tools like MetaGO [15] and the more resource-efficient KmerGO [16] leverage long k-mers (up to 40 bp) to identify group-specific sequences that distinguish between clinical traits or disease states.

The necessity of direct usage on reference genomes can be bypassed by leveraging the ability of Large Language Models (LLMs) to generalize semantic and hidden patterns of biological sequences. This approach allows for the classification of genetic material without the constraints of alignment-based methods. Emerging models like DNABERT-2 [17] and its virus-specific fine-tuned version ViraLM [18] leverage transformer architectures to extract features and distinguish viral signals from complex genomic backgrounds. When genomic sequences are converted into embeddings, they function as coordinates in a high-dimensional latent space. It is hypothesized that sequences from different organism types naturally form distinct clusters within this space based on their intrinsic sequence patterns [19].

Previous results indicate that k-mer spectra can effectively compare datasets and identify novel organisms without a reference genome. Alternatively, a more direct approach measures the read coverage of contigs to determine the relative abundance of specific sequences across datasets. Additionally, sequences can be clustered to identify “exclusive clusters” present in only one dataset, providing a clear metric for detecting unique biological features. Finally, emerging sequences can be detected by tracing changes in the occupancy of the latent space of an LMM.

In this work, we develop and test several different methods to recognise the spreading sequences in metagenomic data using comparative metagenomics. We develop four methodologies: alignment-free k-mer analysis, differential coverage estimation, sequence clustering and tracing changes in latent-space of an LLM.

## Methods

All four methods compare baseline and comparative datasets to detect sequences that are more abundant in the comparative one.

### Coverage-based method

The coverage-based pipeline maps short reads to assemblies from the comparative dataset to detect contigs with higher coverage in the comparative dataset than in the baseline one (Figure 1).

**Figure 1.**
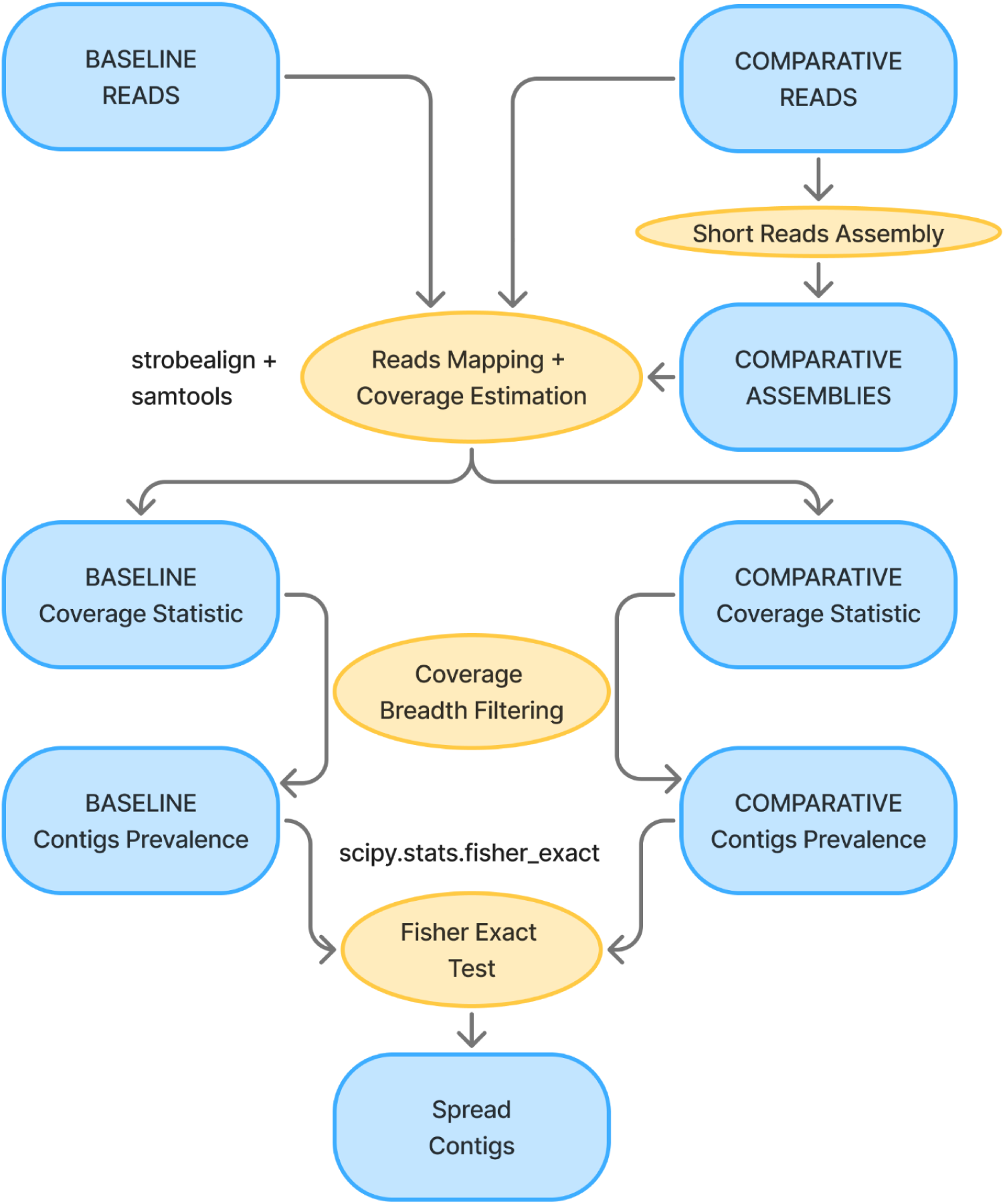
The scheme of coverage-based method.

De novo assembly of short reads is performed using MEGAHIT with the --meta parameter. Contigs shorter than 1000 bp are excluded from the analysis (using the --min-contig-len 1000 option). Only reads from the comparative dataset are used for assembly, based on the hypothesis that the spread sequences are more abundant in these samples.

Read mapping is performed with strobealign [20] with default parameters. The evaluation of coverage breadth is performed with samtools coverage [21].

Contigs are assessed for presence in each sample based on a predefined coverage breadth threshold. A summary table is generated to record the prevalence of each contig across samples from both the baseline and comparative datasets.

Finally, a one-sided Fisher’s exact test is used to evaluate the statistical significance of the difference in contig prevalence between the two datasets. The p-value threshold is adjusted using the Bonferroni correction, where the number of hypotheses corresponds to the total number of contigs. Contigs with a significant difference in prevalence are identified as spread.

### k-mer based method

The k-mer-based pipeline is designed to identify sequences with increasing abundance through a comparative analysis of k-mer prevalence between the baseline and comparative datasets (Figure 2).

**Figure 2.**
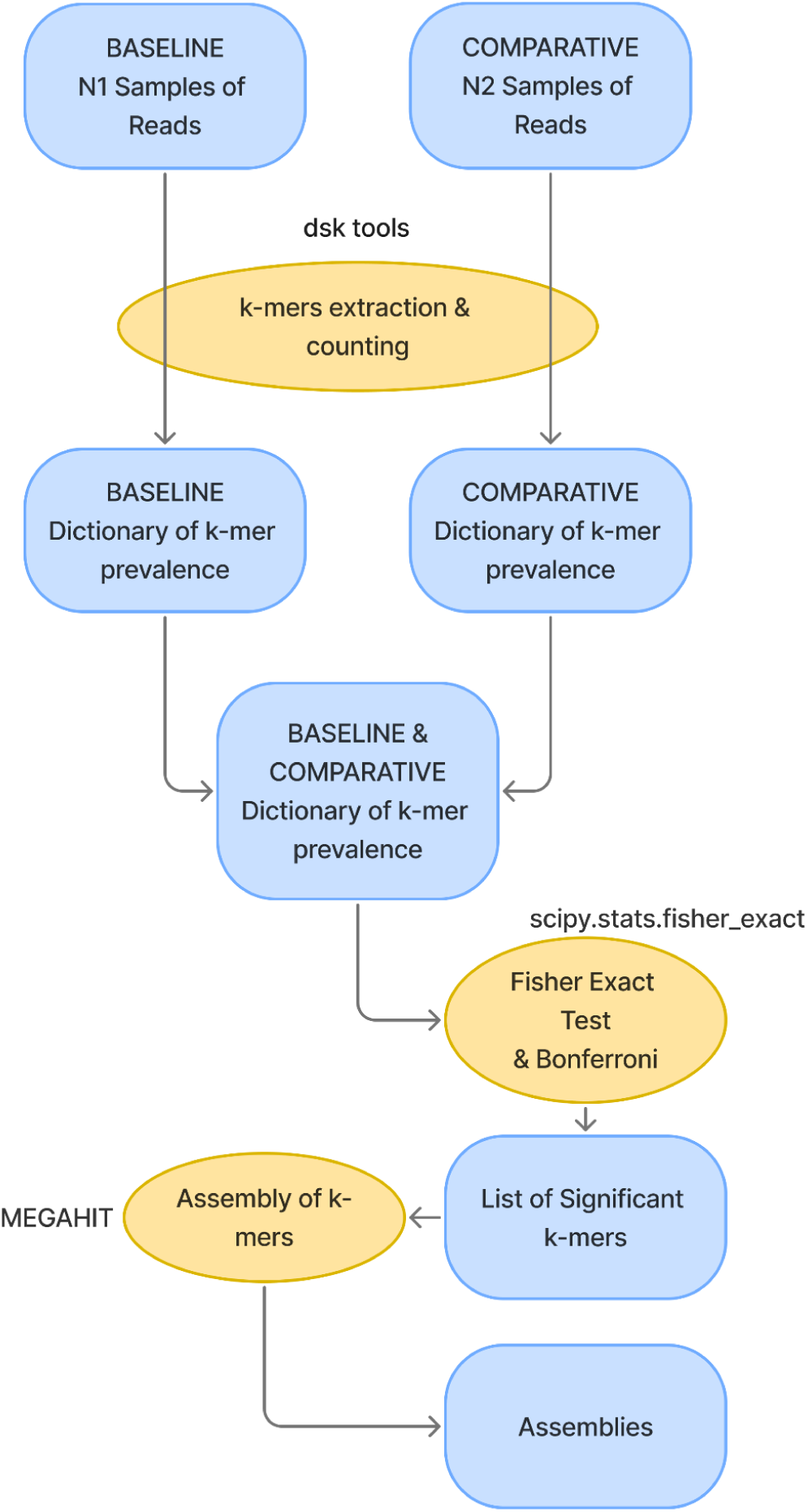
The scheme of the k-mers based method. Blue fields - data, yellow - instruments, green - reference data for measurements of the methods performance

The process begins with k-mer profiling using the DSK tool [22], which calculates the frequency of short sequences in each sample. Subsequently, the dsk2ascii utility converts binary data into text files, which serve as input for specialized Python scripts

During the data aggregation stage, a custom Python script is applied to each dataset. This script encodes nucleotide sequences into integer representation to optimize memory usage. During the processing, the script merges k-mer frequencies across all samples within a dataset and performs primary filtering, removing rare sequences that appear fewer than five times. The results are stored as sorted text files and binary dumps to ensure reproducibility and rapid data access.

The next step is the integration of information from the two datasets. Using an efficient algorithm for merging sorted lists, the program matches profiles between the baseline and comparative datasets. The script compares dictionaries from both datasets and combines them into a single table.

### Statistical evaluation

In the next step, the statistical evaluation of the differences in abundance is provided by Fisher’s Exact Test (fisher_exact from scipy). We process data in large blocks (chunks) to conserve RAM and utilize result caching (lru_cache) to significantly accelerate calculations for repeating frequency values. The primary objective of this stage is to identify sequences whose frequency is statistically significantly higher in the second datasets compared to the first.

The resulting p-values undergo a filtering procedure after the Bonferroni correction for multiple testing. k-mers with significantly increased prevalence are then converted back into nucleotide sequences and assembled de-novo using MEGAHIT.

### Clustering-based method

The clustering-based pipeline first groups sequences by nucleotide identity and then identifies clusters significantly enriched in the second time point.

Initially, each sequence is tagged with its corresponding sample name to maintain metadata throughout the analysis. The full sequence pool is then clustered using MMseqs2 with the parameters --cov-mode 1 and --cluster-mode 2. Following clustering, sequences are partitioned into separate FASTA files based on their cluster assignments. During this stage, clusters containing fewer than three sequences and individual sequences shorter than 1,000 nucleotides are excluded.

A custom Python script subsequently parses these FASTA files to create a dictionary-based data structure. For each cluster, the script records sequence identifiers, lengths, and associated sample names. Using this metadata, the pipeline quantifies the distribution of samples from the baseline and comparative datasets within each cluster.

Finally, the statistical significance of cluster overrepresentation in the comparative dataset is determined using Fisher’s exact test. The p-value threshold is adjusted using the Bonferroni correction, where the number of hypotheses corresponds to the total number of clusters.

### LLM (Large Language Model)-based method

The principle of the LMM pipeline involves converting sequences into coordinates within a multidimensional latent space, followed by the identification of over-occupied regions.

The ViraLM model [18] is employed to transform nucleotide sequences into embeddings. The original source code was modified to extract embeddings from each transformer block using the register_forward_hook function. Because each sequence consists of X tokens, the model yields an output matrix of dimension (X, 768) for each layer. To generate a single representative embedding vector for the entire sequence, token embeddings are averaged across all X tokens using the numpy.mean function, resulting in a vector of dimension (768). These embeddings are extracted from each of the 12 transformer layers.

Subsequently, Principal Component Analysis (PCA) is applied to the data. The PCA model is trained on the embeddings of both datasets (implemented via sklearn.decomposition.PCA with prior scaling using sklearn.preprocessing.StandardScaler) with n_components=2 (Fig. 3). To detect shifts in the distribution of data points within the PCA space, the space is partitioned into isometric bins (chunks), where the boundaries correspond to the global minimum and maximum coordinates of the combined dataset. The value of each chunk represents the number of data points it contains. By default, this process generates two 20×20 occupancy matrices (grid size may vary by experiment), where bins with identical coordinates represent the same spatial regions.

**Figure 3.**
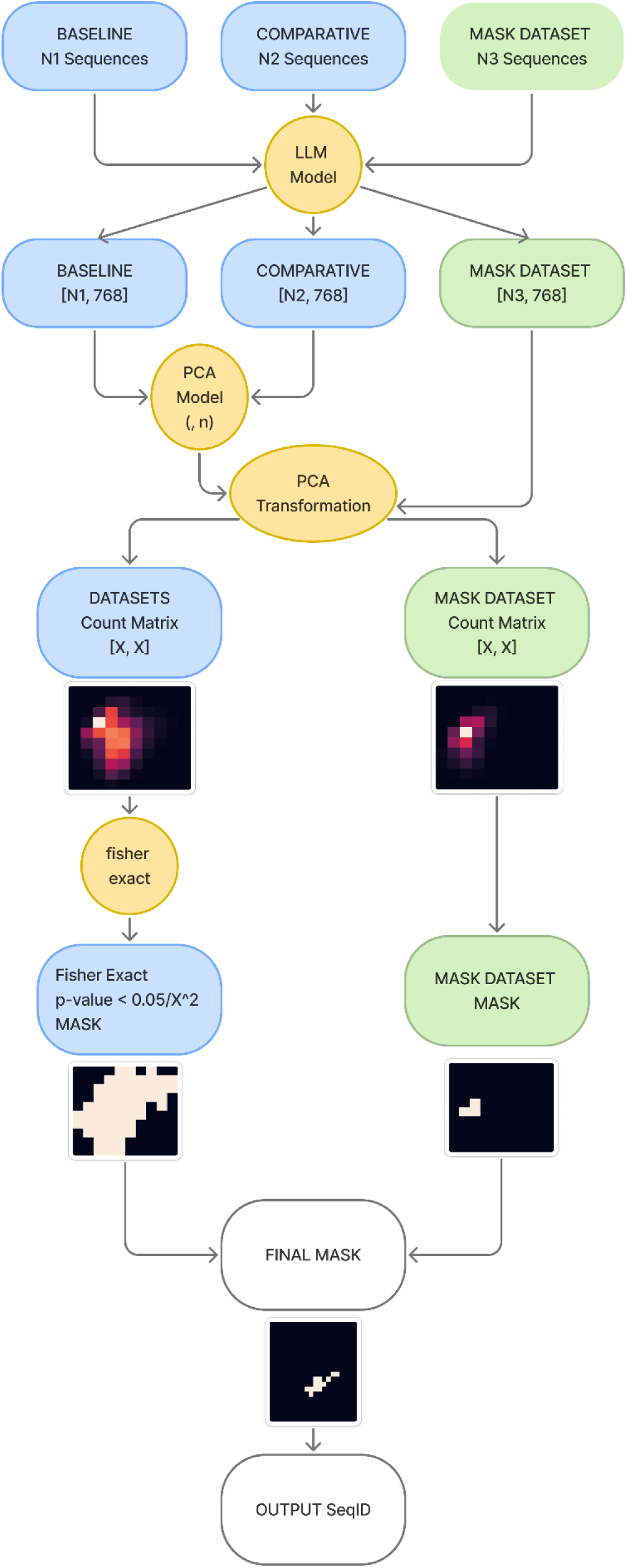
The scheme of LLM-based pipeline.

A mask is generated to identify bins where the values in the comparative dataset exceed those in the baseline dataset, highlighting regions with higher point density. To assess the statistical significance of chunk occupancy, numerical values are compared pairwise using Fisher’s exact test (scipy.stats.fisher_exact). The Bonferroni correction is applied to account for multiple comparisons, with the number of hypotheses defined by the total number of chunks. Finally, a statistical significance mask is constructed based on the adjusted p-values to isolate regions of significant expansion.

To localize specific taxa within the established PCA space, a reference projection is performed using known sequences (e.g., from the ICTV database). These reference sequences are transformed into the same latent space using the pre-trained ViraLM model and the existing PCA parameters. A taxon-specific occupancy mask is then generated to delineate regions associated with the target taxon. This allows for the direct spatial correlation between emerging clusters in the metagenomic data and known biological taxons.

### Datasets

#### Pre/pandemic Datasets collection

We retrieved total RNA-seq datasets of human nasopharyngeal swabs from the NCBI SRA using the following search query: “nasopharyngeal swab, nasopharynx, respiratory OR nasal OR oral NOT gut, Homo Sapiens, Human, RNA-Seq [Strategy]”.

The data were split into pre-pandemic (dated before 2020, n=367, baseline) and post-pandemic (collected after 2020, n=2059, comparative) sets. To ensure a balanced analysis, a subsample of 367 files was randomly selected from the comparative dataset to match the size of the baseline dataset.

The final dataset comprised 734 sequence read runs with a total size of 5,098 GB; the pre-pandemic and post-pandemic timepoints contained 8.86B and 7.43B reads, respectively. After the de novo assembly, a total of 403,403 contigs were generated for downstream analysis.

#### Semi-synthetic dataset creation

To evaluate the performance of the proposed pipelines, a semi-synthetic dataset was constructed.

A set of 200 samples was selected from the post-pandemic collection, which characterized a high degree of homogeneity. These samples were split into two equal groups (n=100) to serve as the baseline and comparative datasets for the semi-synthetic evaluation. Raw sequencing data were retrieved from the EBI ENA database.

For the samples from the comparative dataset, we added the simulated reads of the *Egovirales* MAG giant virus [23]. These reads were generated using InSilicoSeq (v2.0.1) [24], with the proportion of viral reads in each sample determined by an exponential distribution. The total number of viral reads was calculated by multiplying the file’s original read count by the generated percentage. The *Egovirales* MAG giant virus was chosen because it is a newly discovered viral taxon which was not used in ViraLM training.

We tested several values for the λ parameter in exponential distribution (10, 20, 40, 60, 100, 200, and 2000), where λ controls the scale of sampled values: larger λ corresponds to a smaller mean value (1/λ). For each λ and dataset, we calculated the total volume of viral reads added to the second time point of the semi-synthetic data.

#### Runtime benchmark dataset

The benchmark dataset is constructed through an iterative doubling process (10, 20, 40, 80 samples) using synthetic data from two semy-synthetic datasets. The initial 10-sample subset is randomly selected from the first and second timepoints. Each subsequent step expands the dataset by retaining all samples from the previous iteration and adding an equal number of new unique samples (e.g., the 80-sample set comprises the 40 previous samples plus 40 new ones)

#### Calculating performance metrics

To evaluate the sensitivity and accuracy of the methods, an additional validation step was implemented. All the contigs derived from the data (either from the original metagenome assembly or the spread-k-mer assembly) were aligned to the reference genome of the target organism (SARS-CoV-2 in pandemic dataset, Egivirales MAG for synthetic dataset). Specifically, a BLASTN alignment was performed with the following parameters: -outfmt ’6 std qlen qcovs’. The resulting alignments were filtered to retain only high-quality matches with query coverage per subject (qcovs) > 80% and percent identity (pident) > 95%. Contigs meeting these criteria were then used to construct confusion matrices for performance evaluation.

Contigs with increased prevalence were classified as true positives (TP) if they matched the reference genome, and as false positives (FP) otherwise. Conversely, contigs without increased prevalence were classified as false negatives (FN) if BLASTN identified them as belonging to the reference genome, or as true negatives (TN) if they showed no match to the reference.

Precision and recall were calculated to evaluate the performance of the classification. Precision was defined as TP / (TP+FP), while recall was calculated as TP / (TP+FN).

However, these terms were applied exclusively to the semi-synthetic . We did not extend these terms to the pre- and post-pandemic dataset, as it contains numerous metagenomic shifts beyond the emergence of SARS-CoV-2, which fall outside the scope of this validation.

#### Reference genomes

Several reference genomes were utilized to validate the computational pipelines. The NC_045512.2 assembly from GenBank served as the reference for the Wuhan-Hu-1 SARS-CoV-2 genome. To generate the semi-synthetic dataset, the Egovirales sequence ANAN16_1_SAMN03203003_S001_BIN_00000093_scaffold_1 was obtained from the supplementary data of the Egovirales preprint.

## RESULTS

### Pipelines and data

We developed four distinct computational strategies to identify and track spreading sequences within metagenomic datasets. The first is a coverage-based pipeline that assembles contigs from the comparative dataset and maps short reads from both datasets to these contigs to identify sequences exhibiting significantly increased coverage breadth in the comparative dataset. The second approach involves differential k-mer analysis, which compares k-mer frequency spectra between baseline and comparative datasets to identify significantly spread k-mers for subsequent assembly into contigs. The third clustering-based pipeline contigs from both datasets based on nucleotide identity, subsequently flagging clusters that are disproportionately composed of sequences from the comparative dataset. Finally, we implemented an LLM-based pipeline that leverages a Large Language Model to project sequences into high-dimensional embeddings; the pipeline then identifies “over-occupied” regions, defined as areas of the latent space with significantly higher density in the comparative dataset, and classifies all sequences residing within these regions as overrepresented.

To evaluate the performance of our methods in detecting the spread of novel organisms, we compiled a large-scale metagenomic dataset representing real-world viral shifts. This collection consists of 734 human nasopharyngeal RNA-seq runs (5,098 GB), structured as a matched comparison between pre-pandemic (367 samples, 8.86B reads) and post-pandemic (367 samples, 7.43B reads) cohorts. This dual-period structure provides a natural baseline to test the pipelines’ ability to distinguish spread SARS-Cov-2 sequences from stable background microbiota, ultimately yielding 403,403 contigs for analysis.

To further validate these methods in a controlled environment, we constructed a semi-synthetic dataset. This consists of 200 highly homogeneous samples split into baseline and comparative groups (n=100 each), where the latter was enriched with simulated reads of the Egovirales MAG giant virus. Using this novel taxon—which was absent during model training—allowed us to precisely measure detection sensitivity across a gradient of viral concentrations (λ=10 to 2000). Finally, we established a benchmarking framework of iteratively doubling subsets (10, 20, 40, and 80 samples) to evaluate how computational performance scales with increasing data volume.

### Performance of the four methods

#### Performance on the pandemic dataset

##### The four methods were tested on pandemic data

The coverage-based method identified the highest number of contigs as spread, totaling nearly 16,000. Of these, only 800 (approximately 5%) were annotated by Kraken2 as belonging to SARS-CoV-2. The majority of the remaining contigs were of human origin, while a smaller portion was identified as bacterial. While this method achieved 100% coverage width of the SARS-CoV-2 genome, SARS-CoV-2 contigs made up 5.06% of the detected set, and the SARS-CoV-2 contigs accounted for 24.4% total length of the spread contigs (Table 1).

**Table 1.** Results of different methods on pandemic dataset.

|  | Coverage-based | Clustering-based | k-mer-based | ViraLM |
| --- | --- | --- | --- | --- |
| Unclassified | < 1% (30) | < 1% (7) | ~1,5% (1) | 20% (1408) |
| Classified | > 99% (15751) | 99% (6258) | 98,5% (59) | 80% (5887) |
| Bacteria – *Pseudomonadati* | 4.32% (772) | 15% (943) | 10% (6) | 36% (1635) |
| Bacteria – *Bacillati* | 0 | 1.2% (75) | 3.3% (2) | 11.87% (866) |
| Eukaryota – *Homo Sapiens* | 74,76% (11798) | 70% (4387) | 3.3% (2) | 1270 (17.41%) |
| Eukaryota– *Fungi* | 1.22% (192) | < 1% (40) | 1.5% (1) | < 1% (15) |
| Viruses – SARS-CoV-2 | 5.07% (800) | 12.31% (771) | 78% (47) | 24% (1786) |
| Viruses – Other RNA viruses | 0 | 0 | 0 | <1% (10) |
| Viruses - other DNA Viruses | 0 | 0 | 0 | ~1% (91) |

**Table 2.**
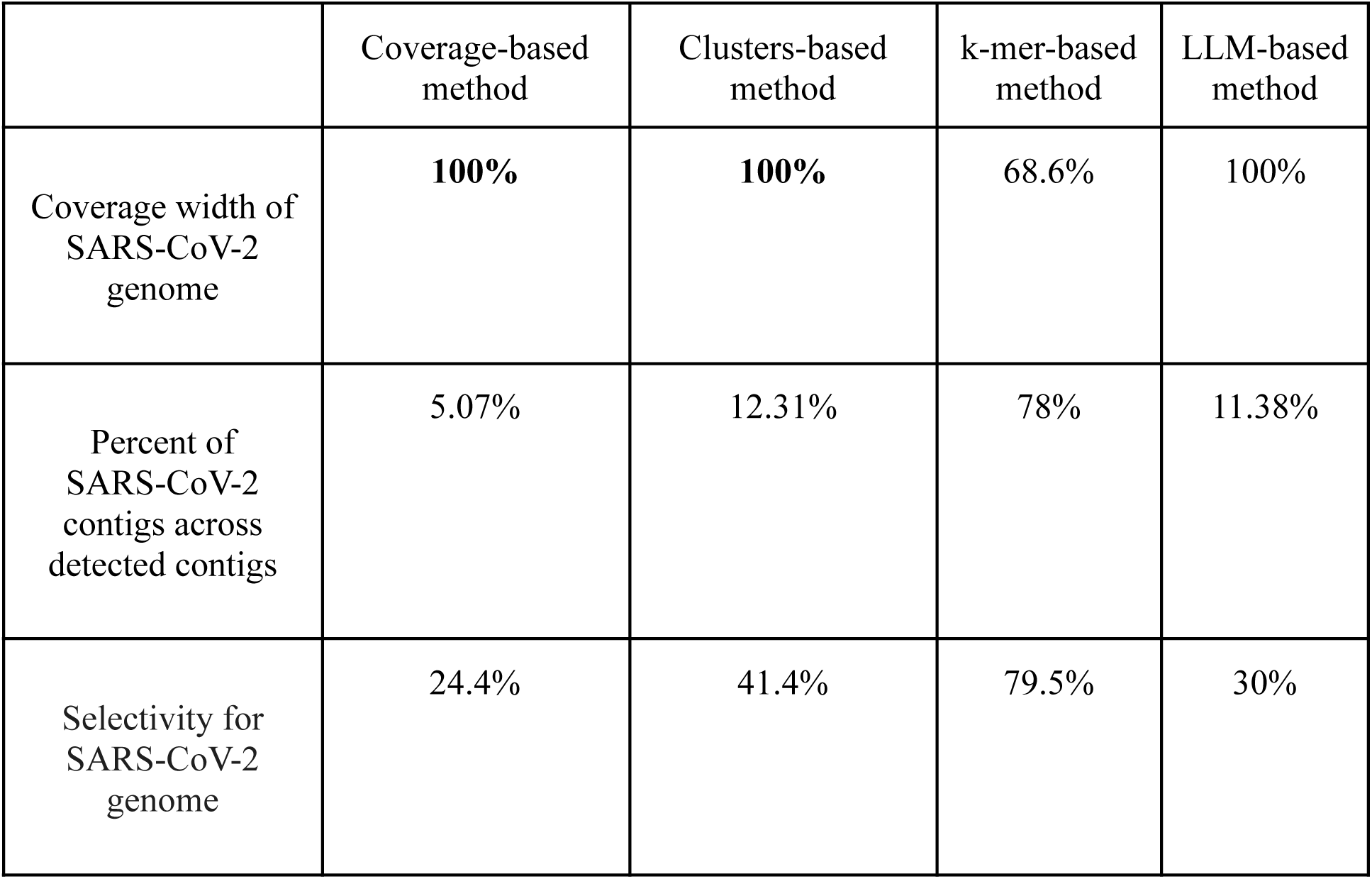
Performance of four methods on the pandemic dataset.

|  | Coverage-based method | Clusters-based method | k-mer-based method | LLM-based method |
| --- | --- | --- | --- | --- |
| Coverage width of SARS-CoV-2 genome | **100%** | **100%** | 68.6% | 100% |
| Percent of SARS-CoV-2 contigs across detected contigs | 5.07% | 12.31% | 78% | 11.38% |
| Selectivity for SARS-CoV-2 genome | 24.4% | 41.4% | 79.5% | 30% |

On the other hand, the clustering-based method yielded a total of 6,265 contigs. Within this set, 771 contigs were identified as belonging to SARS-CoV-2, accounting for 41.4% of the total length of all positive contigs. Notably, these sequences provided 100% coverage of the SARS-CoV-2 genome.

After applying the k-mer pipeline to the pandemic data, contigs were assembled from k-mers with increased representation. A total of 60 contigs were obtained (Table 1). Of these, 47 contigs were identified as belonging to SARS-CoV-2. They have a total length of 20,304 nucleotides and cover more than 60% of the SARS-CoV-2 HU-1 reference genome. The smaller portion of contigs consists of various sequences from humans or cellular microorganisms. This method reached the SARS-CoV-2 genome coverage width of 68.6%. SARS-CoV-2 sequences represented 78% of the detected contigs and 79.5% of the total length of contigs identified by this approach.

In turn, the LLM-based pipeline classified more than 7,000 original metagenomic contigs as spread. SARS-CoV-2 sequences account for approximately 1800 (∼25%) of these contigs. The majority of the remaining sequences consist of various contigs annotated by Kraken2 as bacterial. This LLM-based method achieved 100% coverage width of the SARS-CoV-2 genome. Within the total set of detected sequences, SARS-CoV-2 contigs accounted for 11.38%, while representing 30% of the total length of spread contigs

#### Performance on the semi-synthetic dataset

To evaluate the performance of the four methods at varying concentrations of spreading sequences, we constructed a semi-synthetic dataset to minimize uncontrolled variables, such as batch effects and the presence of other spreading organisms. The dataset is constructed from a homogeneous set of total RNA sequencing data derived from human nasopharyngeal samples, into which Egovirales sequences were introduced. Simulated short reads of Egovirales were incorporated into the samples according to an exponential distribution with varying λ parameters (the larger the λ, the smaller the average value with the mean is equal to 1/λ).

To measure performance, we utilize two primary metrics: the coverage breadth of the Egovirales genome and selectivity, defined as the ratio of the total length of spread Egovirales contigs to the total length of all spread contigs.

**Table 3.** Performance of four methods on the semi-synthetic dataset.

| Amount of egovirales reads | Coverage breadth (contig-based) | Nucleotide selectivity (contig-based) | Coverage breadth (Clustering) | Nucleotide selectivity (clustering) | Coverage breadth (k-mer) | Nucleotide selectivity (k-mer) | Coverage breadth (LLM) | Nucleotide selectivity (LLM) |
| --- | --- | --- | --- | --- | --- | --- | --- | --- |
| 791 K | 99.80% | 89.80% | 99.17% | 99.20% | 99.17% | 100% | 100% | 6.89% |
| 435 K | 99.80% | 85.68% | 99.03% | 98.49% | 98.91% | 100% | 100% | 6.34% |
| 66 K | 98.66% | 88.16% | 99.05% | 98.96% | 98.42% | 100% | 100% | 4.89% |
| 30K | 99.78% | 86.57% | 99% | 99% | 98.30% | 100% | 100% | 2.33% |
| 10 K | 95.84% | 68.30% | 22% | 97% | 0 | - | 100% | 0.07% |
| <1K | 99.55% | 77.78% | 0 | - | 0 | - | 100% | 0.33% |

The comparative analysis of four pipelines demonstrates distinct performance across varying concentrations of spread Egovirales sequences. The contig-based approach proved to be the most balanced, maintaining a coverage breadth of approximately 99% even in low-abundance datasets while keeping nucleotide selectivity above 68% at the 10K read threshold. The clustering-based pipeline performed effectively at higher volumes, but its coverage decreased to 22% at 10K reads (with 97% selectivity) and was unable to produce results at lower concentrations. Similarly, the k-mer-based method provided high selectivity in high-volume datasets but was sensitive to data depletion, with its detection capability reaching zero at the 10K read mark. Finally, the LLM-based pipeline achieved 100% coverage breadth across all tested volumes, though this high sensitivity was accompanied by a lack of precision, as selectivity peaked at 6.89% and fell below 1% in the sparsest datasets.

### Scaling of Computation Time with Sample Size

**Figure 4.**
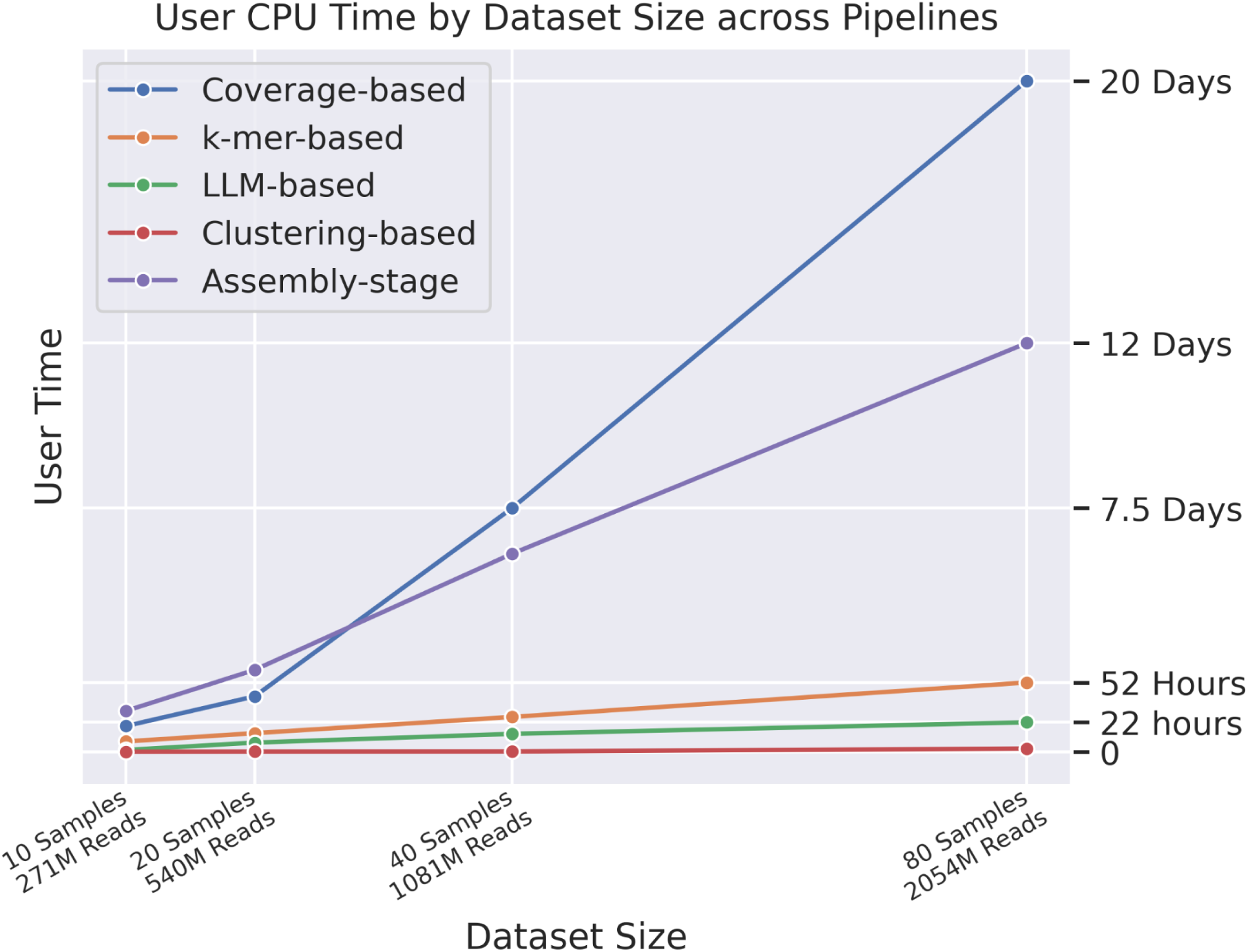
The user computation time across different pipelines. The assembly stage, used in all pipelines except the k-mer-based, is separated into a separate line

Beyond direct performance metrics, computational efficiency remains a critical factor in practical applications. To evaluate this, we developed a series of semi-synthetic datasets of varying sizes and measured the execution time for each approach.

The results demonstrate that while the coverage-based method achieves the highest performance, it is the most resource-intensive. Specifically, on a dataset containing 2B paired-end reads, the computational demand reaches 3M (20 Days) seconds of user CPU time using 20 cores of an Intel(R) Xeon(R) Silver 4210R. In contrast, the k-mer-based method exhibits significantly better efficiency on the same data volume, requiring 39K seconds of real CPU time—nearly a tenfold reduction compared to the coverage-based approach. Meanwhile, the LLM-based pipeline requires 83K seconds of real-time execution using a single CPU thread and GPU (RTX A5000).

## DISCUSSION

”Disease X” represents a hypothetical pathogen with pandemic potential that likely originates and spreads silently in wildlife before spillover to humans. Because viruses often circulate without clear symptoms—similar to the early spread of HIV or TTV —proactive mapping of the wildlife “virosphere” is critical for early detection.

Since viruses lack universal genetic markers, researchers rely on two primary detection strategies: reference-dependent and -independent methods. Since reference-based organism searches rely on homology, they can fail if the new sequence is highly divergent [25]. Alternatively, the spreading sequence may escape researchers’ attention if it is incorrectly taxonomically annotated [26, 27]. On the contrary, methods that do not use homology allow us to identify spreading sequences without prior knowledge of the object.

A relevant example of a dataset capturing the emergence of a novel organism is the metagenomic sequencing of human nasopharyngeal swabs collected before and after the beginning of 2020. Prior to this period, such datasets primarily reflected the commensal microbiota and common respiratory pathogens. Post-2019, a vast majority of nasopharyngeal sequencing research has been dedicated to the detection and genomic characterization of SARS-CoV-2.

We have developed four reference-independent pipelines designed to detect prevalent organisms by comparing two distinct metagenomic datasets, such as those representing different time points. These methods facilitate the identification of emerging or enriched biological entities without reliance on prior genomic knowledge.

The k-mer-based approach decomposes short reads into k-mer sets and generates occurrence tables for each dataset, recording the number of samples containing each k-mer. Statistical comparison then identifies k-mers significantly overrepresented in the second dataset.

The coverage-based method utilizes de novo assembled contigs from the comparative dataset as reference sequences. Short reads from both time points are mapped back to these contigs. Based on defined coverage thresholds, contigs are classified as present or absent across samples, allowing for the isolation of sequences exclusive to the second dataset.

The cluster-based pipeline performs clustering of assemblies from both datasets by nucleotide identity. Each resulting cluster is evaluated for enrichment based on the origin of its contigs, specifically filtering for clusters that consist almost entirely of sequences from the second time point.

Finally, the ViraLM-based approach leverages a large language model to transform contigs into high-dimensional embeddings. These vectors are projected into a latent space where statistical analysis identifies regions with significantly higher density in the second dataset, indicating the presence of divergent or novel viral signals.

To evaluate the performance of our analytical methods, we developed a semi-synthetic dataset using *Egovirales* sequences. The *Egovirales* virus order was specifically selected because its sequences are highly divergent from the training data used for the ViraLM model so the model cannot rely on previously learned patterns to identify these viral elements. This involved using baseline metagenomic samples with adding of varying concentrations of *Egovirales* sequences to quantify both sensitivity and selectivity of our methods.

On pandemic dataset, the k-mer-based approach demonstrated high selectivity and moderate sensitivity, achieving a SARS-CoV-2 genome coverage width of 68.6% and a high nucleotide selectivity of 79.5% on the pandemic dataset. However, its dependence on exact sequence matches suggests potential limitations for rapidly mutating pathogens. High mutation rates or genomic rearrangements may cause contig fragmentation, as the pipeline likely partitions sequences into multiple conserved segments. In semi-synthetic tests, the method’s effectiveness declined significantly once the target sequence count fell below a specific threshold, reaching 0% coverage when target reads accounted for less than 1% of the total dataset.

The coverage-based approach emerged as the most robust in terms of sensitivity, consistently reaching 100% coverage width of the SARS-CoV-2 genome. It maintained superior performance on semi-synthetic data, with nucleotide selectivity remaining above 95% even at low viral concentrations (<1% of all reads). Despite these advantages, the method is significantly limited by its high demand for computational resources; on a dataset of 2B reads, it required 20 days of user CPU time. To improve practical utility, future implementations could incorporate a pre-filtering step to remove the large volume of human sequences (74.76% of detected contigs) prior to mapping, which would reduce the computational burden without sacrificing accuracy.

The clustering-based pipeline achieved full genome coverage and 41% selectivity when evaluated on the pandemic dataset. On semi-synthetic data, the method performed robustly down to an expected genome coverage of 10x. At this level, the pipeline maintained 97% selectivity, although genome coverage decreased to 22%. With less sparse genome coverage, the method loses its effectiveness. This observed decline in coverage is likely attributable to the isolation of genomic segments, which leads to their fragmentation into distinct clusters and a subsequent reduction in cluster representation.

In contrast, the ViraLM-based pipeline showed mixed results. While it achieved 100% genome coverage on the pandemic dataset, its performance on semi-synthetic data was characterized by extremely low selectivity, dropping to 0.07% at λ=200. This lack of precision potentially renders the current LLM-based approach unsuitable for the intended goal. The low selectivity is likely a consequence of the isometric partitioning of the latent space; the appearance of target sequences in a cell causes all sequences in that cell to be labeled positive. While computationally faster than mapping (83K seconds), the trade-off in selectivity remains a critical barrier.

In summary, the k-mer-based approach provides high selectivity but is limited by its reliance on exact matches, making it vulnerable to sequence mutations and low viral concentrations. The coverage-based method offers the highest sensitivity and robustness, consistently achieving complete genome coverage, though it is hindered by extreme computational demands. The clustering-based pipeline performs well at moderate coverage depths, yet it suffers from fragmentation and reduced effectiveness when processing sparse data. Finally, while the ViraLM-based pipeline is the most computationally efficient, it exhibits critically low selectivity, rendering it unreliable due to high false-positive rates caused by its latent space partitioning.

The development of metagenomic sequencing techniques facilitates the continuous monitoring of microorganisms circulating within the biosphere. By analyzing genetic material from wastewater [28], domestic animals [29] wildlife [30, 31], researchers can detect ecological shifts and establish early warning systems for characterizing emerging pathogens.

Given the prevalence of ’dark matter’ in metagenomic datasets [32], reference-independent methods are a valuable addition to homology-based workflows for identifying organisms that lack significant similarity to known genomes.

